# Marine bacterial resistomes integrate ecological adaptation with anthropogenic amplification: genome-resolved insight along a gradient of human impact

**DOI:** 10.64898/2026.03.13.711565

**Authors:** Dmytro Spriahailo, Adenike Adenaya, Thorsten Brinkhoff, Thomas Reinthaler

## Abstract

Antibiotic resistance genes (ARGs) are ubiquitous in marine environments, yet whether their distribution primarily reflects anthropogenic pollution or intrinsic ecological functions remains unresolved. We used genome-resolved metagenomics to characterize resistomes in 371 genomic operational taxonomic units (gOTUs) across a gradient of human impact: the heavily impacted Baltic Sea, the moderately impacted North Sea, and the minimally impacted West Greenland shelf. ARG density was distinctly elevated in the Baltic Sea (3.20 ARGs Mbp^−1^) relative to the North Sea (1.90) and West Greenland (1.67), which did not differ significantly from each other, suggesting a relatively uniform oceanic baseline. Variance partitioning revealed that taxonomic identity explained 20.1% of ARG density variation, with environment contributing 11.4%; critically, Baltic gOTUs carried 35.1% more ARGs than predicted from taxonomy alone, indicating environment-driven enrichment beyond baseline taxonomic carriage. Lifestyle-dependent ARG partitioning between particle-attached and free-living prokaryotes emerged only under anthropogenic pressure: free-living bacteria were enriched in multiple resistance classes in the Baltic Sea but showed no differentiation in West Greenland. Only 0.85% of detected ARGs showed ≥70% amino acid identity to clinically characterized sequences in the CARD database, showing that marine ARGs are highly divergent from clinical resistance determinants. Virulence factor annotations were widespread but weakly coupled with ARG abundance, suggesting independent ecological selection. Our results suggest that marine resistomes integrate an intrinsic baseline of ecological functions with selective enrichment of specific resistance mechanisms under anthropogenic pressure, and that genome-resolved approaches are able to quantify the relative contributions of each.

## Introduction

Antibiotic resistance genes (ARGs) have emerged as a significant environmental concern in marine ecosystems, where anthropogenic inputs of antibiotics and resistant bacteria from wastewater discharge, agricultural runoff, and aquaculture can alter microbial community dynamics and pose public health risks [1, 2]. However, resistance genes are not exclusively products of the anthropogenic era. Functional ARGs have been recovered from 30,000-year-old permafrost [3], isolated cave systems [4], and remote soils [5], demonstrating that many resistance determinants are ancient features of microbial genomes predating clinical antibiotic use by millions of years. Previous research has shown that environmental homologs of clinically relevant resistance mechanisms (including efflux pumps, beta-lactamases, and glycopeptide resistance operons) serve ecological roles in detoxification, cell wall remodeling, and inter-microbial competition [6–8]. Despite the growing awareness, the distribution patterns and ecological functions of ARGs in marine systems remain poorly understood. Most marine ARG studies have focused on bulk water samples or anthropogenically impacted sites [9, 10], leaving the partitioning of ARGs between microbial lifestyles, free-living (FL) versus particle-associated (PA), and across unique microhabitats such as the sea-surface microlayer largely unexplored.

ARGs have been detected in Arctic sea ice [11], Antarctic soils [12], and glacial ice [13], suggesting that resistance genes persist even under minimal anthropogenic antibiotic exposure, yet no study to our current knowledge has systematically compared marine resistomes across a gradient of human impact using genome-resolved approaches capable of linking ARGs to their host organisms.

This raises a fundamental question: how much of the observed resistance reflects an ecological baseline, and how much is enriched by human activity? We define the baseline resistome as the conserved complement of resistance-associated genes maintained by natural selection for ecological functions such as xenobiotic efflux, peptidoglycan remodeling, and competitive interactions [6, 7, 14]. Under anthropogenic stress, specific mechanisms may be selectively enriched through direct antibiotic selection, co-selection by pollutants, or enrichment of taxa carrying higher ARG loads [15, 16]. Distinguishing the baseline from its anthropogenically enriched component requires comparing resistomes across environments that differ primarily in human impact intensity, using approaches that can link resistance genes to specific microbial hosts.

The Baltic Sea and the North Sea represent two heavily impacted marine regions, both shaped by strong anthropogenic stressors [17]. In the brackish Baltic Sea, a large human catchment and limited exchange with the North Atlantic drive high nutrient and contaminant concentrations, primarily originating from wastewater discharge, agriculture, and aquaculture activities [18, 19]. The Baltic Sea consistently ranks among the most impacted marine ecosystems globally in cumulative human impact assessments [20], and multiple resistance-associated genes have been detected in Baltic bacterial communities [21, 22]. Compared to the Baltic Sea, the North Sea experiences stronger tidal flushing by Atlantic waters yet remains subject to inputs from major rivers (Rhine, Elbe, and Maas) and coastal industrial and municipal discharges. ARGs detected in the English Channel [23] and in German harbors [24] reflect anthropogenic pressures characteristic of industrialized coastal zones. Within these regions, the sea surface microlayer (SML), the uppermost 1–1000 µm of the ocean surface [25], represents a microhabitat where anthropogenic pollutants [26–28], and physical stressors such as UV radiation [29] converge, potentially intensifying selection for resistance genes. In contrast, waters off West Greenland experience minimal direct pollution, however, climate-driven changes are emerging as key factors shaping microbial communities [30, 31] and may similarly influence resistome composition. These three regions thus span a gradient of cumulative human impact, providing a natural comparative framework for disentangling baseline resistance from anthropogenic enrichment.

Whether ARGs and virulence-factor-associated genes (VFs) co-occur in marine bacteria through co-selection on shared mobile genetic elements or reflect independent responses to distinct ecological pressures remains unresolved [32–35]. Addressing this question is critical for determining whether elevated antibiotic resistance also implies elevated pathogenic potential in marine microbial communities.

Here, we use genome-resolved metagenomics to: (1) characterize resistome composition in metagenome-assembled genomes (MAGs) from three marine regions differing in cumulative human impact; (2) quantify whether ARG prevalence and diversity are enriched above the baseline resistome in anthropogenically impacted environments; (3) examine how ARG distribution differs between microbial lifestyles and microhabitats; and (4) assess the relationship between ARGs and VF-annotated genes across these environments.

## Materials and methods

### Sampling and datasets

Samples were collected from the Baltic Sea (near Rockneby, Sweden), the North Sea (off Helgoland, Germany), and offshore West Greenland (Fig. S1). Baltic Sea samples were collected in May 2021 from surface slicks and non-slick areas as described by Rahlff et al. [36]; the SML was sampled using the glass plate method [37] and the ULW at ~70 cm depth. Seawater was filtered through 5 µm and 0.2 µm polycarbonate filters to separate PA and FL fractions; DNA extraction and sequencing were performed by Rahlff et al. [36] and we reanalyzed their raw sequences. North Sea SML and ULW sampling was conducted in September 2022 using a remotely controlled catamaran with a rotating glass disk assembly [38]; ULW was collected at 1 m depth. Samples were filtered through 3 µm, 0.8 µm, and 0.2 µm polycarbonate filters. West Greenland samples were collected in July 2021 using Niskin bottles from surface (~4 m), deep chlorophyll maximum (25–40 m), and near-bottom (60–550 m) depths, then filtered as for the North Sea. Environmental metadata (temperature, salinity, wind speed, nutrients) are provided in Table S4.

The PA fractionation boundary differed between datasets (5 µm in the Baltic vs. 3 µm in the North Sea and West Greenland). This difference is conservative for our main findings: the higher threshold in the Baltic may misclassify some particle-associated bacteria as free-living, potentially underestimating Baltic PA-associated ARG density. In total, we analyzed 77 metagenomes: 8 from the Baltic Sea, 24 from the North Sea, and 45 from West Greenland.

### DNA extraction, sequencing, MAG recovery, and gOTU detection

For North Sea and West Greenland samples, DNA was extracted using the DNeasy PowerWater Kit (Qiagen) and sequenced on the Illumina NovaSeq S4 PE150 XP platform at the Vienna BioCenter Core Facilities. Read-level taxonomic profiling used SingleM v0.18 [39]. Quality control and assembly followed a Snakemake v9.13 [40] workflow with parameters listed in Table S1. Three separate co-assemblies were generated using MEGAHIT v1.2 [41] (assembly metrics in Table S2). Binning employed MetaBAT2 v2.15 [42], SemiBin2 v2.2 [43], GenomeFace [44], and VAMB v5.0 [45], with graph-based refinement (GraphBin2 v1.3 [46]) and consolidation (Binette v1.1 [47]). Bins were dereplicated at 99.5% ANI (dRep v3.6 [48]), manually refined in anvi’o v8 [49] targeting those with ≥90% completeness and ≥5% redundancy (CheckM2 v1.1 [50]). Full pipeline source code is available at https://github.com/sprdmt/bass-fastq-to-mag.

Finally, the MAGs were dereplicated again at 95% ANI to define gOTUs. gOTU presence per sample required ≥50% read-mapping breadth, ensuring that a majority of the representative genome was covered and reducing false positives from spurious mapping to conserved regions [51]. After filtering, 66 of 77 metagenomes retained detectable gOTUs. Of 2,699 ARG-encoding genes, 2,695 in 348 gOTUs passed filtering.

### ARG and VF annotation

MAGs were classified using GTDB-Tk v2.4 [52]. Genes predicted by Prodigal v2.6 [53] were annotated for ARGs using DeepARG v2.0 [54] (≥50% identity, ≥0.8 probability), AMRFinderPlus v4.0 [55] (≥70% identity), CARD v6.0 [56] (≥70% identity), and KOfamScan [57] (e-value ≤1e−5, resistance-associated KO terms). For genes detected by multiple tools, a fixed priority hierarchy (DeepARG > AMRFinderPlus > CARD > KOfamScan) retained a single annotation per gene (sensitivity analysis using six alternative orderings produced 99.8–100% concordance). Resistance class labels were standardized; “unclassified”, “heavy metal”, and “antiseptic” categories were excluded. Virulence factors were identified by DIAMOND blastp against the virulence factor database (VFDB) [58] (e-value ≤1e^−5^, ≥50% identity, ≥50% query coverage).

### Statistical analysis

ARG prevalence comparisons used Fisher’s exact tests with FDR correction. ARG density (ARGs Mbp^−1^) was compared using Kruskal–Wallis and pairwise Wilcoxon tests. Differential abundance of ARG classes was tested in edgeR v4.2 [59] using negative binomial models with non-ARG exposure offsets and TMM normalization. Post-hoc power analysis used the edgeR exactTest framework to estimate, for each comparison, the minimum log_2_ fold change that our design could detect with 80% statistical power. This indicates an 80% probability of identifying a true effect of that magnitude significant at α = 0.05, given the observed dispersion and sample sizes. Robustness was assessed by bootstrap resampling (10,000 iterations), permutation tests (9,999 iterations), and leave-one-out jackknife analysis. Variance partitioning used linear models with environment and bacterial class as predictors. Environmental correlates were tested using Spearman correlations and multi-variable linear models with nested F-tests. ARG–VF associations were modeled using GAMs with negative binomial distributions and genome-size offsets, with VF annotation sensitivity tested at 50%, 60%, and 70% identity thresholds. All analyses used R v4.5 [60]. Detailed statistical and bioinformatic methodology is provided in the Supplementary Methods.

## Results

### Prokaryotic community structure from read-based taxonomic annotation

Microbial community composition was primarily structured by lifestyle (FL vs. PA), with region as a secondary factor (Fig. 1). FL communities were consistently dominated by Pelagibacteraceae (SAR11), while PA communities were dominated by Flavobacteriaceae and Rhodobacteraceae. West Greenland showed pronounced depth-related shifts, including dominant Nitrosopumilaceae (Archaea) in the FL fraction at pelagic depths. Alpha diversity, estimated by the Shannon index, varied significantly across regions (*p* < 0.01). Microbial fraction explained 47.5% of Shannon diversity variance, with region and depth each contributing ~15%. Family-level richness differed significantly among regions (*p* < 0.001), with region explaining 55.9% of variance.

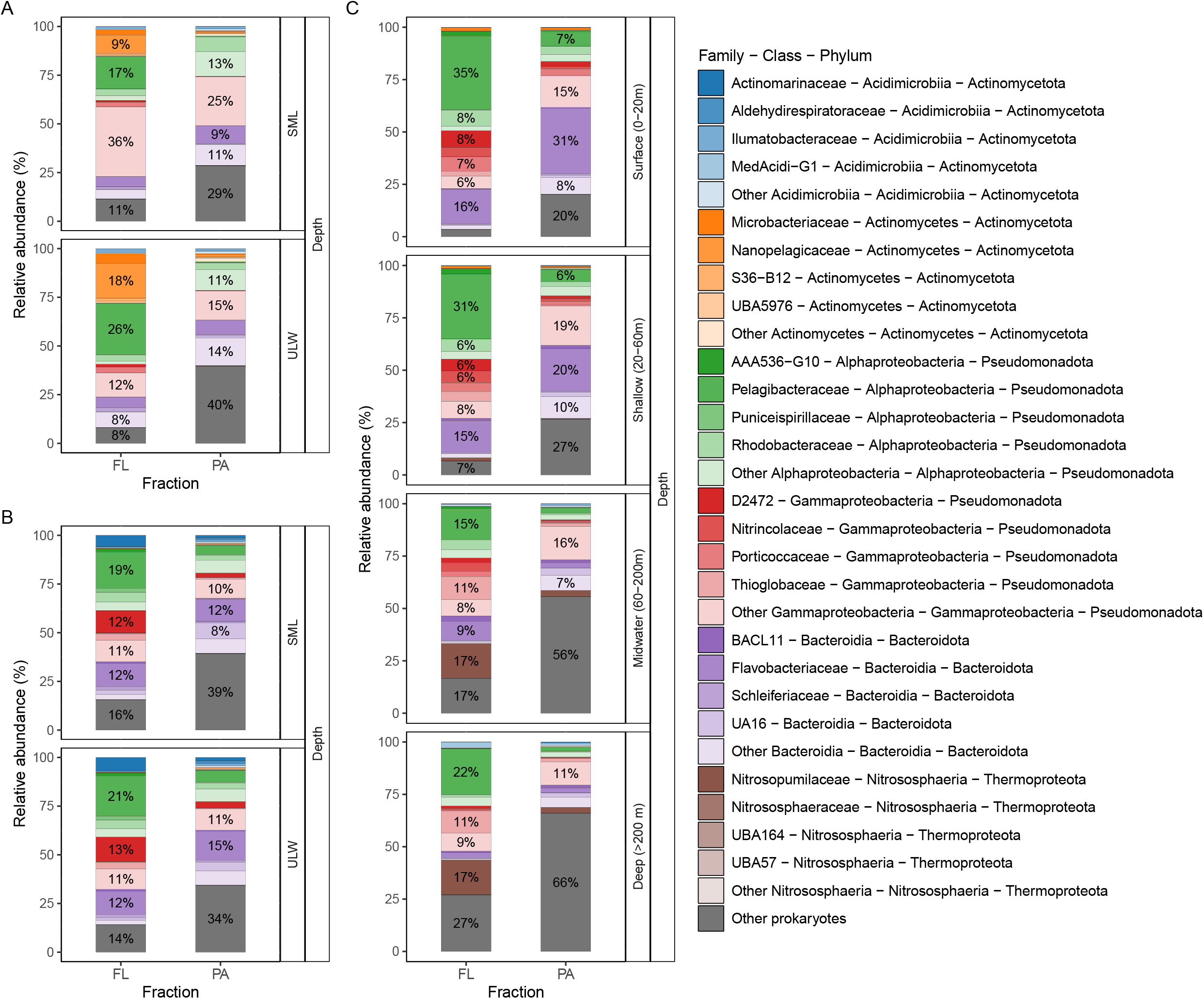

### Taxonomy and distribution of gOTUs

A total of 371 MAGs were recovered: 80 from the Baltic Sea, 172 from the North Sea, and 119 from West Greenland. These MAGs had a median completeness of 96.6% and a median redundancy of 1.6% (CheckM2). Clustering at ≥95% average nucleotide identity yielded 371 gOTUs, including 176 putative novel species lacking close GTDB-Tk matches. Based on read mapping (≥50% breadth), 89, 185, and 135 gOTUs were detected in the Baltic Sea, North Sea, and West Greenland, respectively. Only 5 gOTUs were shared among all regions, whereas most were site-specific (72, 156, and 104 unique gOTUs, respectively) (Fig. 2A).

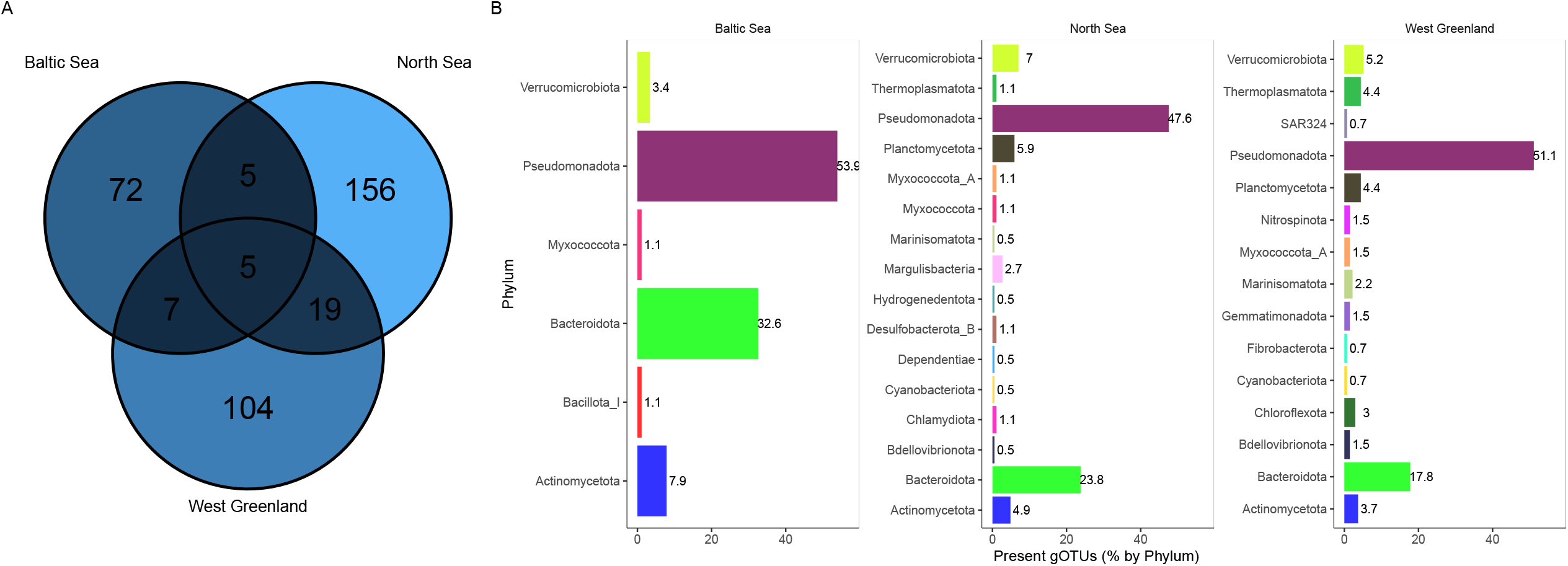

At the phylum level, Pseudomonadota comprised approximately 50% of gOTUs across all sites, followed by Bacteroidota and Actinomycetota (Fig. 2B). Minor fractions included Verrucomicrobiota, Myxococcota, and Thermoplasmatota, while SAR324, Nitrospinota, and Chloroflexota occurred only in West Greenland.

### Abundance of resistance genes in gOTUs

The annotation pipeline identified 2,699 unique ARG-encoding genes across 15 resistance classes in 348 gOTUs, with KOfamScan contributing 70.6% of annotations and DeepARG 28.6%. Only 23 (0.85%) of identified ARG homologs showed ≥70% amino acid identity to the Comprehensive Antibiotic Resistance Database (CARD) sequences, indicating that the vast majority of marine ARGs are highly divergent from clinical counterparts.

Most gOTUs carried at least one, mainly efflux-associated multidrug resistance genes (Baltic Sea 92%, North Sea 78%, West Greenland 84%; Fig. 3A, 3B). The prevalence of specific resistance classes differed by region: glycopeptide, macrolide-lincosamide-streptogramin (MLS), tetracycline, β-lactam, and aminoglycoside resistance genes occurred in 64–39% of Baltic gOTUs, 52–24% in the North Sea, and 47–23% in West Greenland (Fig. 3C and Table S3). The most prevalent individual ARGs were efflux-associated *tolC* (53–66%), *acrA* (19–54%), *acrB* (18–54%), and *macB* (20–33%), with consistently elevated prevalence in the Baltic Sea (Fig. 3C and Table S3).

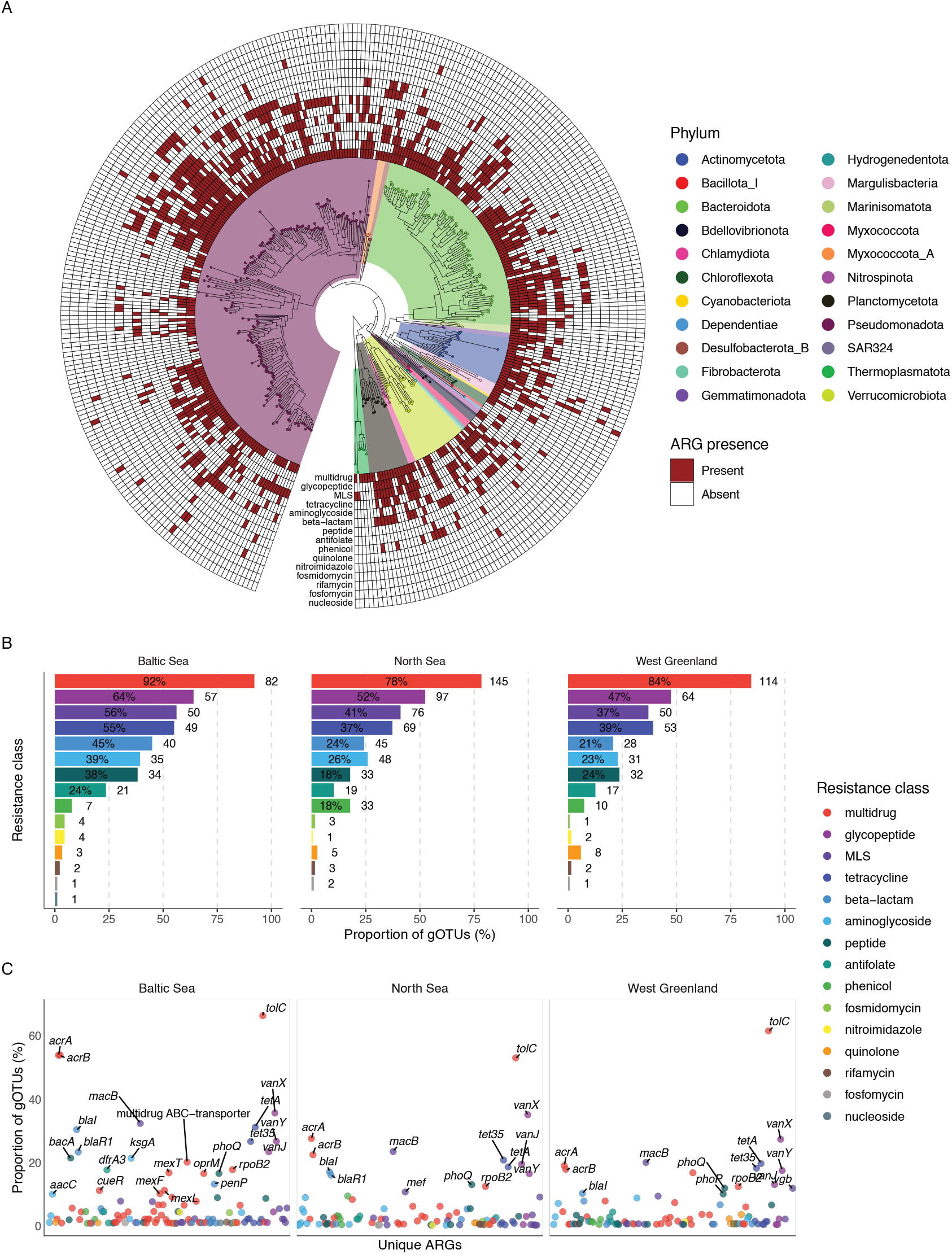

Across taxa (pooled across regions; Wilcoxon one-vs-rest, FDR <0.05), ARG densities (ARGs Mbp^−1^) showed taxon-specific patterns with substantial heterogeneity. At the phylum level, Pseudomonadota (n = 171 gOTUs) exhibited weak overall enrichment (Cliff’s δ = 0.21), however, masking within-phylum variation: Gammaproteobacteria (n = 85) showed moderate-to-strong enrichment (δ = 0.45) and Actinomycetes (n = 6) – strong enrichment (δ = 0.68). Lower ARG densities characterized Planctomycetota (δ = −0.37) and Thermoplasmatota (specifically, Poseidoniia: δ = −0.80) at the phylum level, with Planctomycetia (δ = −0.34) and Alphaproteobacteria (δ = −0.17, n = 86) at the class level. At the family level, Alteromonadaceae (median 5.87 ARGs Mbp^−1^, n = 8) and Burkholderiaceae (4.45, n = 6) showed the highest, while Thalassarchaeaceae (0.66, n = 5) and Rhodobacteraceae (1.32, n = 15) the lowest ARG densities. Genome sizes did not differ significantly among environments (Kruskal–Wallis, *p* = 0.81, n = 371), indicating that observed ARG density differences are not confounded by genome size. A linear model accounting for both environment and genome size confirmed that environment explained substantially more variance in ARG density than genome size (environment: F = 38.4, *p* < 0.001; genome size: F = 6.4, *p* < 0.05; R^2^ = 0.18).

### Habitat and lifestyle-associated ARG enrichment

The Baltic Sea showed significant enrichment in 9 of 15 ARG classes relative to West Greenland (log_2_FC >1, FDR <0.05; Fig. 4A). Despite our sample size imbalanced dataset (Baltic Sea n = 8, West Greenland n = 34) the edgeR post-hoc power analysis indicated that only 4 out of 9 ARG classes had a |log_2_FC| smaller than ≈ 2.4 and fell below the <80% power threshold. Comparing the North Sea (n = 24) and West Greenland showed similar but weaker trends with a minimum detectable |log_2_FC| ≈ 1.6 at 80% power. Within the Baltic Sea, SML samples were enriched in nitroimidazole, quinolone, and antifolate resistance relative to ULW (Fig. 4B), with all three showing large size effects (|log_2_FC| > 4, n = 4 for SML and ULW). Between lifestyles, FL communities in the Baltic Sea showed higher proportions across most ARG classes than PA communities, while the North Sea showed a more variable pattern (Fig. 4C). No significant differences between PA and FL communities were observed in West Greenland.

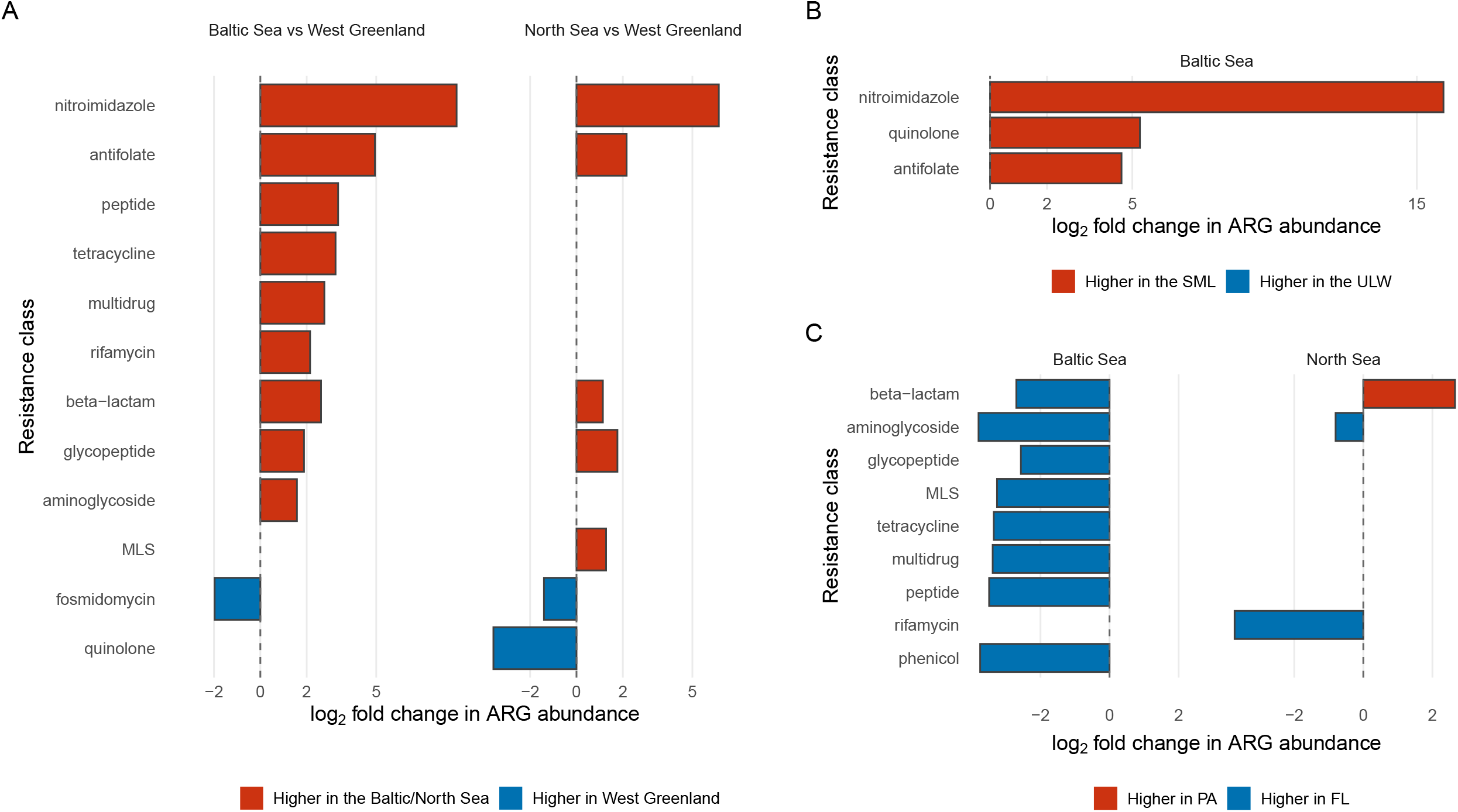

Variance partitioning of gOTU-level ARG density (total ARGs per genome size, n = 384 gOTU-sample observations across all three regions) revealed that taxonomy (bacterial class) explained 20.1% of total variance, while environment explained 11.4%, with 5.2% shared between the two factors (additive model R^2^ = 0.37; ANOVA: environment F = 46.1, *p* < 0.001; class F = 3.8, *p* < 0.001).

Within-taxon tests confirmed that environmental effects persisted after accounting for taxonomic composition. Among the classes represented in all three environments (n ≥ 3 per environment), ARG density differed significantly within Bacteroidia (Kruskal– Wallis *p* < 0.001, n = 93), Gammaproteobacteria (*p* < 0.001, n = 90), and Alphaproteobacteria (*p* < 0.05, n = 99), with Baltic Sea gOTUs consistently showing the highest median densities. The expected ARG density based on taxonomic composition alone (weighted by the global class mean) was 2.59 genes Mbp^−1^ for the Baltic Sea, 2.24 for the North Sea, and 2.29 for West Greenland. Observed densities exceeded expectations by 35.1% in the Baltic Sea but fell 8.9% and 16.0% below expectations in the North Sea and West Greenland, respectively, suggesting enrichment driven by environmental factors rather than taxonomy alone.

### Environmental predictors of ARG density

To identify physicochemical drivers of resistome variation, per-sample mean ARG density (averaged across present gOTUs) was correlated with environmental variables (Table S4). Salinity was the strongest single predictor of per-sample ARG density in the North Sea (Spearman ρ = −0.601, *p* < 0.001, n = 66), with the correlation also significant within West Greenland, where salinity varied with depth and water mass (ρ = −0.785, p < 0.001). Baltic Sea salinity was invariant (6.6 PSU) and could not be tested within-region. Additionally, within West Greenland, ARG density correlated positively with dissolved oxygen and chlorophyll a, and negatively with nitrate, phosphate, silicate, and water density, linking ARG carriage to the photic surface layer. A multivariate linear model including temperature and salinity showed that environmental factors accounted for the majority of per-sample variance in ARG density (R^2^ = 0.788), and environment remained significant even after accounting for temperature and salinity (F = 5.2, p = 0.008), indicating that region-specific factors beyond measured physicochemical gradients contribute to resistome differentiation.

### Relationship between ARGs and virulence-factor genes

Among the 371 gOTUs, 368 (99.2%) encoded at least one ARG or VF, and 328 (88.4%) contained virulence-associated genes. VF counts ranged from 3 to 294 per genome, with 13 gOTUs exceeding 100. The most frequent virulence-related functions were linked to adherence, motility, nutritional/metabolic factors, stress survival, and biofilm formation.

Baltic Sea gOTUs exhibited the highest median ARG densities (3.20 ARGs Mbp^−1^; n = 88), followed by the North Sea (1.90 ARGs Mbp^−1^; n = 170) and West Greenland (1.67 ARGs Mbp^−1^; n = 126). ARG densities were significantly higher in the Baltic than in both the North Sea and West Greenland (*p* < 0.001), while North Sea and West Greenland did not differ significantly (*p* = 0.17) (Fig. 5A). VF densities followed similar regional patterns (Fig. 5B), with the Baltic significantly elevated. Bootstrap resampling (10,000 iterations) confirmed non-overlapping 95% confidence intervals for median ARG density across all three regions, and leave-one-out analysis confirmed that no single Baltic gOTU drives the observed elevation.

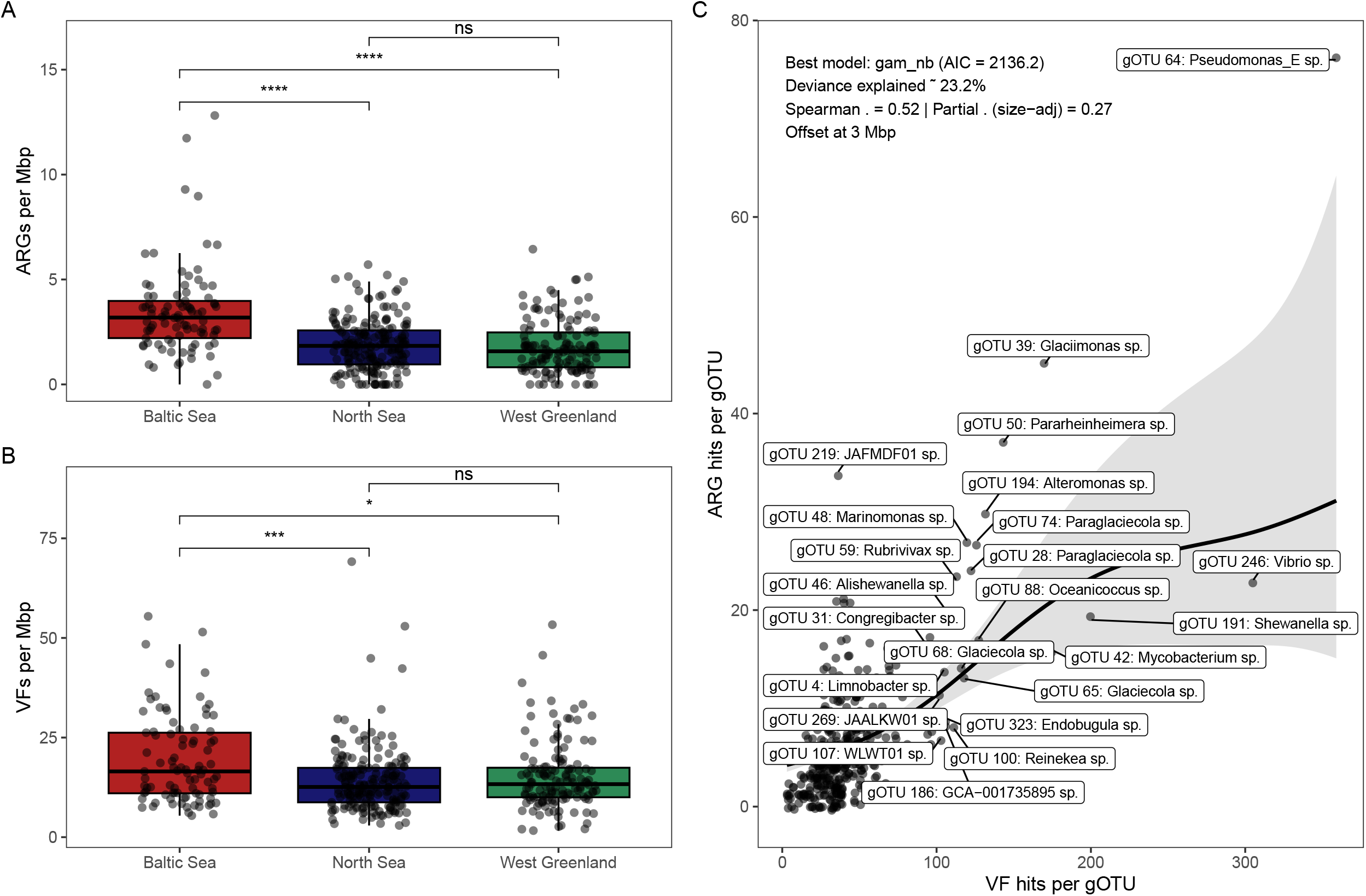

A generalized additive model with a negative-binomial distribution and genome-size offset revealed a significant non-linear association between VF and ARG counts (deviance explained = 23.2%; Spearman ρ = 0.52; partial ρ = 0.27, *p* < 0.001; n = 371; Fig. 5C). The ARG–VF correlation was also robust to stricter VF annotation thresholds (Spearman ρ = 0.40 at 60% identity/coverage; ρ = 0.24 at 70% identity), indicating that the co-occurrence is not an artefact of permissive database matching.

## Discussion

### Ecological structuring of microbial communities

Bacterial community composition was primarily structured by lifestyle, with region and depth as secondary factors, consistent with well-described partitioning by substrate availability [61, 62]. The dominance of Pelagibacteraceae (SAR11) in the FL communities [63] reflects their adaptation to oligotrophic conditions and efficiency in utilizing low-molecular-weight substrates [64]. In contrast, PA bacterial communities were dominated by known copiotrophic polymer-degraders, including Flavobacteriaceae and Planctomycetota, reflecting their specialization for processing particle-associated high-molecular-weight organic matter [65].

The elevated diversity of the Baltic Sea communities likely reflects the overlap between strictly marine and freshwater or brackish taxa together with nutrient enrichment from riverine and anthropogenic sources, which together generate multiple ecological niches [66, 67]. The North Sea showed intermediate alpha diversity, consistent with its dynamic and well-mixed character, whereas the cold, oligotrophic West Greenland waters had lower alpha diversity and a distinct archaeal signal dominated by Nitrosopumilaceae. These patterns reflect the salinity, temperature, and nutrient gradients characteristic of high-latitude coastal ecosystems [67, 68]. The low ARG densities in Alphaproteobacteria (particularly SAR11) likely reflect their oligotrophic lifestyle and streamlined genomes [64], where metabolic efficiency favors reduced gene content, while the higher ARG loads in Gammaproteobacteria may represent a copiotrophic strategy in which diverse detoxification systems enable exploitation of nutrient-rich but chemically variable particle microenvironments. This structuring of communities with lifestyle and along regional gradients was mirrored in the distribution of ARGs.

### Lifestyle and regional patterns in ARG distribution

Total ARG abundance was highest in the Baltic Sea, and lowest in West Greenland, but the distribution between lifestyles varied by environment (Fig. 4C). In the Baltic Sea, FL communities overall showed consistently higher proportions of resistance genes across ARG classes compared to PA fractions, however, the limited sample sizes precluded statistical resolution of individual class-level differences. This pattern of enrichment in dispersed prokaryotes may reflect greater exposure to dissolved selective compounds, including antibiotics and heavy metals, from riverine and wastewater inputs [69], a higher proportion (Fig. 1) of ARG-dense Gammaproteobacteria [70] in FL communities, and reduced diffusion of dissolved compounds into particle microenvironments [71].

The North Sea showed fewer lifestyle-associated differences, and no significant PA– FL partitioning was detected in West Greenland, suggesting that ARGs in this low-impact environment serve conserved physiological functions (e.g., efflux and cell wall biosynthesis) rather than reflecting selective enrichment under anthropogenic pressure. Three ARG classes were significantly enriched in the Baltic Sea SML relative to ULW, all with large effect sizes (Fig. 4B), potentially reflecting elevated pollutant concentrations at the air-sea interface [72], wet and dry atmospheric deposition of aerosols and contaminants [73, 74], or UV-induced oxidative stress [29]. The absence of depth differences in the North Sea and West Greenland, however, suggests effective vertical mixing in these systems. These patterns suggest that bacterial lifestyle and regional environmental characteristics, other than depth, are the primary drivers of ARG distribution in our marine systems. The enrichment of specific ARG classes in Baltic and North Sea gOTUs can be linked to documented anthropogenic inputs. Nitroimidazole resistance enrichment in the Baltic Sea SML likely reflect pharmaceutical contamination, as nitroimidazoles are widely used in human and veterinary medicine and their residues have been detected alongside sulfonamides, tetracyclines, and β-lactams in European wastewater effluents and Baltic coastal waters [75, 76]. Antifolate resistance enrichment is consistent with the documented presence of sulfonamides, which are among the most frequently detected antibiotics in Baltic waters [21], while tetracycline and β-lactam enrichment aligns with regional antibiotic use patterns [21, 77]. This selective enrichment of resistance mechanisms, rather than the general elevation of all ARGs, may provide a more sensitive indicator of anthropogenic impact in marine monitoring programs.

Baltic Sea ARG densities were distinctly elevated above both the North Sea and West Greenland (Fig. 5A), with the latter two not differing significantly from each other. This pattern is independently supported by the Halpern Cumulative Human Impact Index (CHI), which ranks the Baltic Sea among the most impacted marine ecosystems globally (CHI > 8), well above the North Sea (CHI ≈ 4–6) and West Greenland (CHI < 2) [20]. The widespread occurrence of ARGs in minimally impacted West Greenland waters suggests that a substantial fraction of resistance-annotated genes serve ecological functions beyond antibiotic defense, constituting a baseline resistome upon which anthropogenic pressures selectively act. Interpreting West Greenland as a pristine reference nonetheless requires caution, as cold temperatures, glacial meltwater inputs, and low nutrient availability independently shape its resistome. While variance partitioning confirms that environment remains a significant predictor of ARG density after accounting for taxonomic composition, West Greenland is more accurately characterized as a complex low-impact environment rather than a simple pristine baseline.

### Possible ecological roles of resistance-annotated genes

The widespread distribution of ARGs in our results supports the emerging view that many resistance-annotated genes function as an ecological toolkit for detoxification, structural maintenance, and stress response [6, 7]. Efflux and transport systems dominated the ARG annotations, with multidrug, glycopeptide, MLS, tetracycline, β-lactam, and aminoglycoside categories occurring in decreasing order of prevalence. Among these, *tolC* and functionally related *acrA* and *acrB* were among the most frequent, consistent with the ubiquity of resistance–nodulation–division efflux systems in marine and soil bacteria (Fig. 3B, 3C) [78, 79]. Rather than representing antibiotic-specific adaptations, these complexes likely evolved to export metabolic byproducts and xenobiotics [14, 80], maintaining intracellular homeostasis under variable organic matter and pollutant loads [81]. In eutrophic surface layers, they may alleviate metabolic stress from accumulated allelopathic compounds, reactive oxygen species, and competition for limiting inorganic nutrients [82, 83]. While both glycopeptide and β-lactam resistance genes were widely prevalent across regions, they may serve primary roles in peptidoglycan biosynthesis [84, 85] rather than antibiotic defense. Both systems modify peptidoglycan precursors, a critical function in marine environments where salinity fluctuations impose continuous osmotic stress requiring dynamic cell wall adjustments [86–88]. Although vancomycin resistance was initially characterized in clinical *Enterococcus* [89] strains, and penicillin-binding proteins were first described in pathogenic contexts, their homologs occur widely in environmental bacteria [90–92], suggesting ancient genetic modules maintained for structural homeostasis. Some bacteria can even utilize β-lactams as carbon sources [93], further supporting ecological rather than defensive functions.

Importantly, this baseline can still be modulated by anthropogenic inputs. The higher ARG abundance in the Baltic and North Sea suggests selective enrichment under nutrient and contaminant pressure [15, 16]. Less than 1% of ARGs in our study (23 of 2,699) showed ≥70% amino acid identity to well-characterized clinical resistance genes in the CARD database, while the vast majority were detected by DeepARG or KOfamScan, indicating substantial sequence divergence from clinical counterparts. This supports the interpretation that most marine ARGs represent environmental homologs with ecological functions rather than recently acquired clinical resistance genes.

### Virulence factor distribution and potential for co-selection

The virulence factor database (VFDB) classifies genes based on their roles in pathogenesis, but in environmental organisms, these genes predominantly encode core ecological functions such as nutrient acquisition, surface colonization or stress tolerance [34, 94]. We therefore interpret VF annotations in our marine genomes as ecological traits rather than indicators of pathogenic potential.

ARGs and VF-annotated genes were near-universal (99.2% and 88.4% of gOTUs, respectively; Fig. 5) suggesting they are standard genomic components. Notably, the most VF-rich lineages were not necessarily the most ARG-rich, indicating that these traits may respond to distinct selective pressures. VF density was slightly elevated in West Greenland (13.3 VFs Mbp^−1^) compared to the North Sea (12.6 VFs Mbp^−1^, Fig. 5B), possibly reflecting the ecological demands of the oligotrophic sub-Arctic environment, where traits such as iron acquisition and motility are essential for survival [95, 96].

The non-linear ARG-VF relationship suggests that bacteria balance investment in resistance and ecological functions (Fig. 5C), with the observed saturation likely reflecting metabolic constraints and fitness trade-offs [33, 97, 98]. The co-elevation of ARG and VF densities in Baltic Sea gOTUs points to environmental stress as a selective driver of broad functional repertoires rather than pathogenic potential [33]. Our results are consistent with the interpretation that resistance genes and VF-annotated genes represent ecological adaptations that pathogens have secondarily co-opted. Whether ARG–VF co-occurrence reflects co-transfer on shared mobile elements or independent ecological selection nonetheless remains unresolved, as plasmids and other mobile genetic elements are poorly recovered by short-read MAG assembly [99].

Taken together, our findings indicate that marine bacterial resistomes are shaped by a combination of conserved ecological functions and anthropogenic selective pressures. The Baltic Sea showed distinctly elevated ARG densities, lifestyle-dependent resistance patterns, and enrichment of specific resistance classes linked to documented pollutant inputs. The North Sea and West Greenland, by contrast, showed similar resistome profiles, suggesting a relatively uniform oceanic baseline. Genome-resolved metagenomics was essential for separating the effects of taxonomic composition from genuine within-taxon enrichment, which is not possible with gene-centric approaches. The predominance of efflux and cell-wall-associated ARG categories, combined with very low sequence similarity to clinical databases, indicates that most marine ARGs likely serve ecological roles. As Arctic and sub-Arctic waters face increasing pressure from shipping, resource extraction, and climate-driven change, the baseline resistome characterized here for West Greenland provides a reference point against which future anthropogenic and climatic impacts can be assessed.

## Supporting information

Figure S1. Geographic sampling locations.

Figure S2. Alpha diversity patterns across environments and fractions.

## Acknowledgements

We thank the captains and crews of R/V Dana and R/V Heincke for their support at sea. We are grateful to Oliver Wurl for initiating and coordinating the BASS research group, and to the Halobates Team for SML sampling. We thank Janina Rahlff for providing the Baltic dataset, and Mathias Wolterink and Carola Lehners for assistance with cruise material transport. We acknowledge the LISC team for maintaining the computer cluster at UBB.

## Supplementary material

Supplementary material is available online.

## Data availability

Raw metagenomic sequences are available at the European Nucleotide Archive (ENA) at EMBL-EBI under BioProject accessions PRJEB104698 (North Sea samples), PRJEB105207 (West Greenland samples), and PRJEB105209 (Baltic Sea samples, reanalysis of BioProject PRJNA855638). All MAGs used for analysis are deposited at ENA under accessions SAMEA120807296–SAMEA120807467 (172 MAGs, North Sea), SAMEA120814707–SAMEA120814827 (121 MAGs, West Greenland), and SAMEA120815998–SAMEA120816077 (80 MAGs, Baltic Sea). All data are accessible via the ENA browser at https://www.ebi.ac.uk/ena/browser/. The analysis script in R is available on Zenodo [100].

