## Supplementary figures and images for "Marine bacterial resistomes integrate ecological adaptation with anthropogenic amplification: genome-resolved insight along a gradient of human impact"

### Figure S1. Geographic sampling locations.

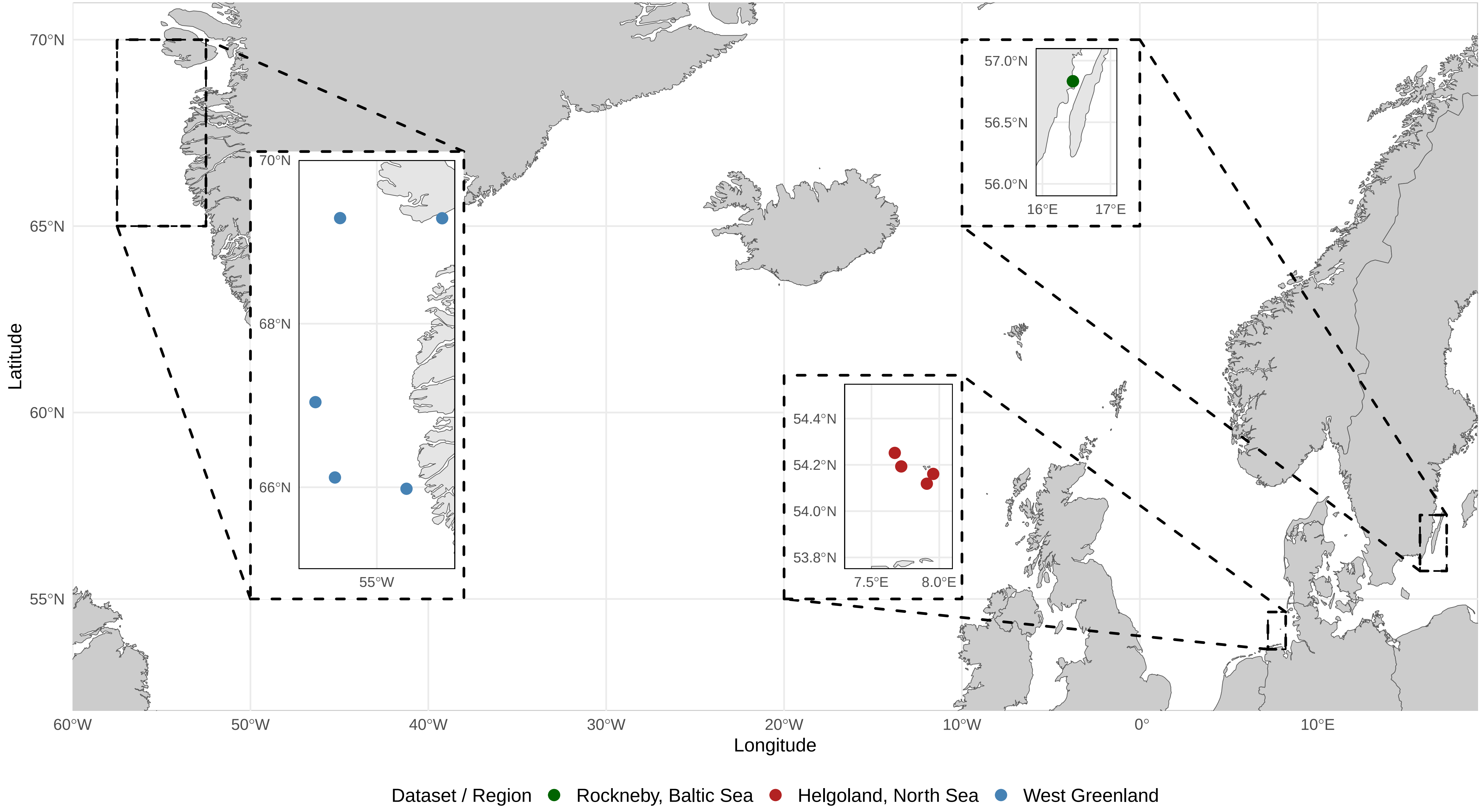

### Figure S2. Alpha diversity patterns across environments and fractions.

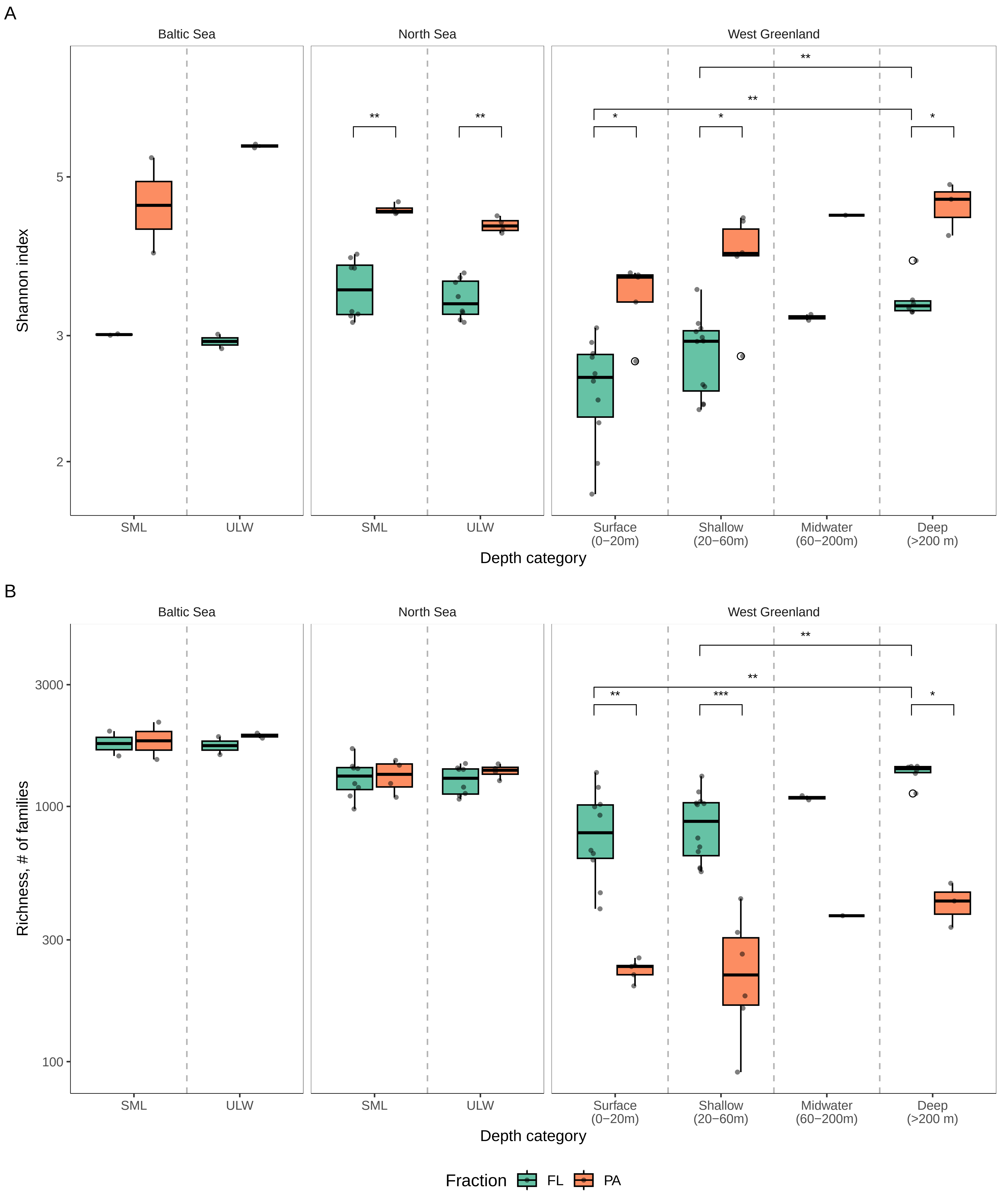
