## Supplementary material for "Marine bacterial resistomes integrate ecological adaptation with anthropogenic amplification: genome-resolved insight along a gradient of human impact": S1 Supplementary Methods.

This document contains detailed statistical and bioinformatic methodology supporting the main text. Section numbering corresponds to the condensed Methods in the main manuscript.

### Read processing and quality control

Quality control of raw reads was performed using FastQC v0.11 [1], followed by adapter trimming with Cutadapt v5.2 [2], trimming of low-quality bases, 5’ trimming on both reads, and discarding reads <50 bp.

### Assembly and binning pipeline details

All quality-filtered reads were pooled within each dataset prior to assembly. Three separate co-assemblies were generated using MEGAHIT v1.2 [3] with k-mer sizes ranging from 29 to 127 (step size 10); assembly metrics are provided in Table S2. Read mapping was performed using Bowtie2 v2.5 [4]. Contig-level coverage profiles (mean depth and related metrics) were calculated with CoverM v0.7 [5]. Initial binning was performed using MetaBAT2 v2.15 [6], SemiBin2 v2.2 [7], GenomeFace [8], and VAMB v5.0 [9] on contigs of 1000 bases or longer. For each binner, contig-to-bin assignments were refined utilizing the assembly graphs using GraphBin2 v1.3 [10]. Consolidation of the refined outputs was performed with Binette v1.1 [11]. Those bins were then dereplicated with dRep v3.6 [12] at 99.5% average nucleotide identity and imported into anvi’o v8 [13] for manual refinement. It targeted bins with ≥90% estimated completeness and ≥5% estimated redundancy in either CheckM2 v1.1 [14] or anvi’o, after which another dereplication at 95% ANI was applied to finalize the genomic operational taxonomic unit (gOTU)-level representative set used for downstream analyses. The precise parameters used in the pipeline can be found in the source code on https://github.com/sprdmt/bass-fastq-to-mag.

### ARG annotation priority scheme

Where multiple tools reported ARG hits for the same predicted gene, we applied a per-gene priority scheme. After tool-specific quality filtering (DeepARG probability ≥0.8 and identity ≥50%; AMRFinderPlus and CARD identity ≥70%), all annotations were merged by gene identifier, and a single annotation was retained for each gene according to the fixed hierarchy. Thus, if a gene was detected by DeepARG, the DeepARG prediction was always used, regardless of whether other tools also called that gene; if DeepARG was absent but AMRFinderPlus and CARD both reported hits, the AMRFinderPlus annotation was retained, and CARD was only used when neither DeepARG nor AMRFinderPlus reported that gene. KOfamScan-derived KO terms were used as a last option when specialised ARG tools did not report a hit for a given gene. The retained record provided both gene-level annotation and resistance class, after standardising resistance class labels and discarding the “unclassified”, “heavy metal”, and “antiseptic” categories.

### gOTU definition and presence gating

MAG dereplication followed a two-step approach. First, near-identical MAGs (>99.5% average nucleotide identity, ANI) were consolidated using dRep v3.4 to remove strain-level redundancy arising from co-assembly of closely related sequences: a standard denoising step that reduces computational burden without losing species-level diversity [15]. After manual inspection and curation of borderline-quality MAGs using anvi’o v8, a second dereplication at 95% ANI defined species-level gOTUs, consistent with the widely accepted species boundary for prokaryotes [16]. For each 95% ANI cluster, we retained a single representative and mapped the gene-level annotations to gOTUs through the representative MAG.

gOTU presence per sample was defined from read-mapping breadth: a gOTU was considered present if it exhibited breadth ≥0.5 (50%) in that sample. The 50% breadth threshold ensures that the majority of a representative genome is covered by mapped reads, reducing false-positive presence calls that may arise from spurious mapping to conserved regions (e.g., rRNA operons or universal single-copy genes). This threshold balances sensitivity (retaining genuinely present low-abundance gOTUs) against specificity (excluding gOTUs detected only through shared conserved sequences). Similar breadth thresholds have been applied in large-scale metagenomic studies [17]. After applying this threshold, 66 of 77 metagenomes (8 Baltic, 24 North, 34 West Greenland) contained detectable gOTUs and were retained for downstream analysis. Of the 2,699 ARG-encoding genes identified, 2,695 genes in 348 gOTUs passed breadth-of-coverage filtering. Unless noted, prevalence and abundance summaries were restricted to present gOTUs. For rate-model analyses, non-ARG background per sample was computed as total mapped counts to present gOTUs minus counts assigned to ARGs; the log of the background served as the exposure offset in edgeR.

### Differential abundance testing

ARG prevalence per resistance class was calculated as the proportion of gOTUs (breadth ≥50%) containing at least one gene from that class, with 95% confidence intervals using Wilson’s method. Between-environment differences were tested using Fisher’s exact tests for each resistance class (15 classes × 3 pairwise comparisons = 45 tests), with Benjamini–Hochberg false discovery rate (FDR) correction at α = 0.05.

ARG density (ARGs Mbp⁻¹) was calculated by dividing total ARG-encoding genes by genome size (from CheckM2). Environment-level comparisons (n = 371 gOTUs) used Kruskal–Wallis tests followed by pairwise Wilcoxon tests with FDR correction. Linear models were fitted to test whether environmental differences in ARG density persisted after adjusting for genome size.

ARG abundances between different sample groups were compared using edgeR v4.2 [18] and limma [19]. Differential abundance of ARG classes was tested in edgeR using a negative binomial generalised linear model with an offset for non-ARG exposure (log of non-ARG read counts from present gOTUs). This offset accounts for sequencing depth while testing for proportional ARG enrichment. The sequencing depths were comparable but uneven across datasets: Baltic Sea samples yielded 167.4 ± 37.0 million read pairs, North Sea samples 100.1 ± 15.2 million read pairs, and West Greenland samples 66.7 ± 37.9 million read pairs. However, the edgeR framework for differential abundance testing uses TMM normalisation to account for differences in library size across samples. Test significance was assessed at FDR <0.05, effect sizes reported as log₂ fold-changes normalised by total non-ARG reads.

### Post-hoc power analysis

Post-hoc power analysis for differential abundance tests was performed using the edgeR exactTest framework. For each significant resistance class, we calculated the achieved power given the observed effect size, dispersion, and sample sizes. Minimum detectable effect sizes at 80% power were calculated for each pairwise comparison.

### Bootstrap, permutation, and jackknife analyses

To assess robustness of ARG density patterns to sampling variability, given the limited number of samples in the Baltic Sea dataset, we performed bootstrap resampling (10,000 iterations, sampling gOTUs with replacement within each environment) and calculated 95% confidence intervals for median ARG density. Permutation tests (9,999 iterations, randomly reassigning environment labels) assessed the significance of pairwise differences. Leave-one-out jackknife analysis evaluated sensitivity to individual gOTUs by iteratively removing each gOTU from the Baltic Sea dataset and recalculating median ARG density.

### Variance partitioning

To distinguish the relative contributions of environment and taxonomy to ARG density variation, we performed variance partitioning using linear models. We fit four models to gOTU-level ARG density data: (1) environment only, (2) bacterial class only, (3) both predictors, and (4) both predictors plus their interaction. Unique and shared variance components were calculated using partial R² values.

To test whether environmental effects persist within taxonomic groups, we performed Kruskal–Wallis tests for ARG density differences among environments, separately for each bacterial class present in all three regions (n ≥ 3 gOTUs per environment). To quantify environment-driven amplification, we calculated expected ARG densities for each environment based on its taxonomic composition (weighted mean of global class-level medians) and compared those to the observed values.

### Environmental correlates of ARG density

To identify the physicochemical drivers of resistome variation, per-sample mean ARG density (averaged across gOTUs present in each sample) was correlated with environmental variables using Spearman rank correlation. Variables tested included temperature, salinity, wind speed, dissolved oxygen, nitrate, phosphate, silicate, chlorophyll a, and water density. Correlations were calculated both pooled across all samples and separately within each region.

Multi-variable linear models were constructed to partition variance in per-sample ARG density: Model 1 included environment (categorical: Baltic, North Sea, West Greenland); Model 2 added temperature and salinity (continuous); Model 3 was the full model. F-tests compared nested models to assess whether the environment influence remained significant after accounting for physicochemical gradients.

### ARG–VF correlation modelling

ARG–VF correlations were modelled using generalised additive models with Poisson or negative binomial distributions and genome-size offsets. Model selection used AIC, with negative binomial providing the best fit. Analyses used R v4.5 (tidyverse, data.table, mgcv, MASS). To assess the sensitivity of ARG–VF correlations to annotation stringency, we repeated VF identification at three identity/coverage thresholds: ≥50% (primary), ≥60%, and ≥70%. Spearman correlations between ARG and VF counts were recalculated at each threshold.
